# Concurrent processing of the prosodic hierarchy is supported by cortical entrainment and phase-amplitude coupling

**DOI:** 10.1101/2024.01.22.576636

**Authors:** Chantal Oderbolz, Sebastian Sauppe, Martin Meyer

**Author notes:** These authors contributed equally and share senior authorship (Sebastian Sauppe, Martin Meyer).

## Abstract

Models of phonology posit a hierarchy of prosodic units that are relatively independent from syntactic structure, requiring its own parsing. Surprisingly, it remains unexplored how this prosodic hierarchy is represented in the brain. We investigated this foundational question by means of an EEG study. Thirty young adults listened to German sentences containing manipulations at different levels of the prosodic hierarchy. Evaluating speech-to-brain cortical entrainment and phase-amplitude coupling revealed that prosody’s hierarchical structure is maintained at the neural level. The faithfulness of this tracking varied as a function of the hierarchy’s degree of intactness as well as systematic inter-individual differences in audio-motor synchronization. The results underscore the role of complex oscillatory mechanisms in configuring the continuous and hierarchical nature of the speech signal and firmly situate prosody as a structure indispensable from theoretical perspectives on spoken language comprehension in the brain.

## Introduction

The speech signal contains hierarchically organized acoustic information. As a consequence of cochlear processing ^1,2^, what becomes available of this signal to the auditory cortex are amplitude modulations (AM) at different frequencies ^3,4^. Most pronounced are amplitude modulations at around 4-8 Hz, but energy also accumulates at the lower modulation rates of ∼2 and ∼1 Hz. In turn, these acoustic features correspond to syllables ^5^, stress or feet ^6^, and intonation phrases ^7^. Phonological models capture these modulations, positing that the speech signal is linguistically organized into units of increasing sizes. These suprasegmental units (spanning over multiple speech sounds) form a prosodic hierarchy that ranges from syllables to intonation phrases ^8,9^. Larger units such as feet are composed of and dominate smaller units such as syllables (see **Figure 1**). Importantly, these models emphasize that prosody “is a complex grammatical structure that must be parsed in its own right” ^8^. Indeed, the information encoded in the prosodic hierarchy provides the listener with cues crucial for creating phonological ^10^, semantic ^11^, syntactic ^12^, and pragmatic ^13^ representations of an utterance. Thus, by extension, generating or comprehending speech requires that the brain efficiently and concurrently process the amplitude modulations associated with the levels of the prosodic hierarchy. Nevertheless, despite accruing evidence for prosody’s significance for language comprehension, this conjecture remains at best sparsely explored. We address this gap in our understanding by means of an EEG study using a simple design involving spoken sentences with prosodic manipulations.

**Figure 1.**
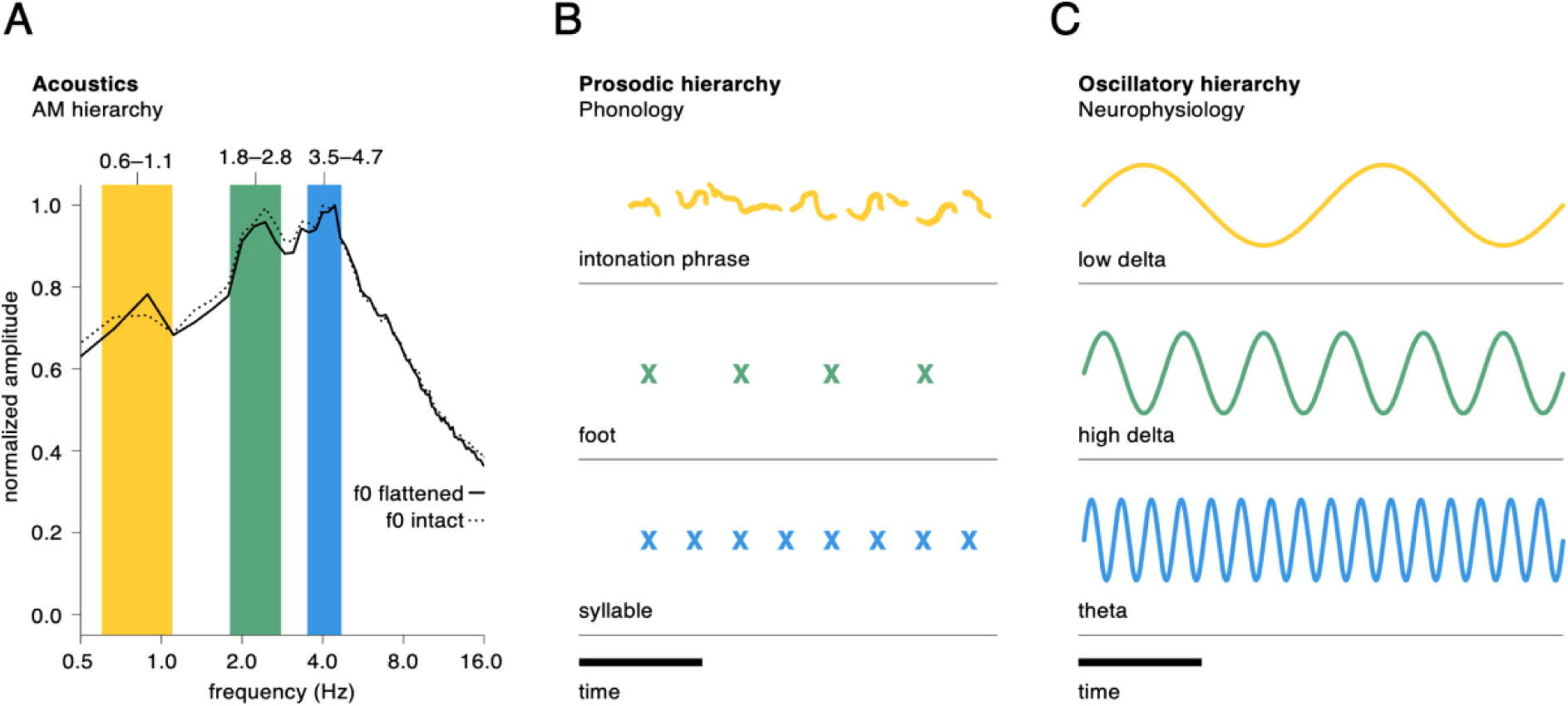
Theoretical framework that forms the basis of the current study. **A** The speech signal accumulates energy at certain modulation frequencies shown here in a plot of the temporal modulation spectrum (yellow: 0.6-1.1 Hz; green: 1.8-2.8 Hz; blue: 3.5-4.7 Hz). **B** Phonological models posit that prosody, like syntax, is hierarchically structured ^9,29,30^. This so-called prosodic hierarchy describes prosodic units and their relationships as relatively independent from syntax and thus as a structure that must be “parsed in its own right” ^8^. The specific prosodic units that constitute the prosodic hierarchy vary slightly from model to model ^31^ with consensus at the very top (i.e., concerning constituents that are defined by a coherent intonation contour) and at the very bottom (concerning constituents within a lexical word) ^32^. Here, we mainly subscribe to Beckman and colleagues’ work ^8,9^ since it is acoustically motivated. Nonetheless, prosody interacts with syntax in important ways ^33^, but this interaction is beyond the scope of this paper. **C** Recent models propose a role for multi-frequency (nested) oscillations in dealing with the hierarchical and continuous nature of the speech signal.

Recent neurophysiological and computational models invoke oscillatory mechanisms through which hierarchical information (such as prosody) can be extracted from continuous speech. On the one hand, the brain employs multi-frequency cortical oscillations that align with the quasi-periodic temporal features of the speech signal (e.g., amplitude modulations) in a process called cortical entrainment ^14–16^. The cyclical nature of cortical oscillations creates temporal windows through which the speech signal is segmented into chunks. Temporal prediction is a natural result of these cycles wherein oscillatory phase is shifted to remain approximately constant across time, allowing for optimal alignment with and segmentation of the speech signal. On the other hand, cross-frequency coupling (CFC) is a phenomenon ubiquitous in dynamic (neural) systems and different architectures of cross-frequency interactions between oscillations are thought to give rise to different functions ^14,17,18^. One putative function linked to phase-amplitude coupling (PAC), or nesting (a form of CFC), is the temporal parsing of continuous stimuli like speech. PAC coordinates the timing and organization of neural activity between the phase of slower and the amplitude of faster oscillations, helping to identify and group relevant acoustic and linguistic units in the speech signal ^19^. Here, our goal was to investigate how the low-frequency amplitude modulations associated with the prosodic hierarchy are concurrently represented in the brain and whether this representation is underpinned by multi-frequency cortical entrainment and a nested hierarchy of oscillations.

Participants listened to short, semantically neutral sentences that, in order to amplify the brain’s ability to concurrently represent different levels of the prosodic hierarchy, contained manipulations (see **Methods**). Each sentence consisted of 16 syllables. At the foot level, stressed and unstressed syllables alternated such that sentences could either follow a predictable regular or irregular (but still legal) stress pattern (e.g., “**Chris**ta **soll**te **da**mals **ein**en **run**den **gel**ben **Kä**fer **ma**len”). At the intonation phrase level, sentences either had a natural or flattened pitch contour ^20^. Based on these sentences’ temporal modulation spectrum, we evaluated cortical entrainment to the syllable/theta (3.5-4.7 Hz), foot/high delta (1.8-2.8 Hz), and intonation/low delta (0.6-1.1 Hz) levels; further, we quantified PAC as a function of prosodic manipulations. Additionally, we examined one more factor. Results from previous studies suggest that the mechanism of cortical entrainment is unequally pronounced across listeners ^21^ with listeners belonging to either a group of high or low audio-motor synchronizers. High synchronizers are characterized by increased cortical entrainment during passive listening to speech and a learning advantage in a phonological word-learning task compared to low synchronizers ^21^. In consideration of these observations, we grouped participants into high (N = 11) and low synchronizers (N = 19) *post hoc*, according to their performance on an established behavioral task ^22^ (see **Methods**).

We predicted the brain to phase-align to each level of the prosodic hierarchy concurrently, such that the influence of prosodic manipulations would be most apparent at the corresponding neural level (i.e., the influence of feet in the high delta and of intonation in the low delta range). However, considering the theoretical framework of the prosodic hierarchy, with larger units linguistically dominating smaller units, we likewise expected a top-down influence of prosodic manipulations on cortical entrainment at lower levels. In this vein, the effect of manipulations at the level of the foot should also be found neurally in the theta frequency range. Specifically, we hypothesized a facilitative effect of rhythmic regularity, manifesting as increased phase-alignment, compared to rhythmic irregularity. Rhythmic regularity, in both speech and auditory sequences, provides cues regarding both the meaningful grouping of units and their timing ^23–25^ — features that the brain can take advantage of to anticipate the occurrence of units in the incoming auditory (speech) stream, resulting in optimal alignment of neural oscillations ^14^. Similarly, the effect of manipulations at the level of the intonation phrase should also be found at subordinate levels. Specifically, we predicted that a naturally varying intonation contour would coincide with increased phase-alignment compared to a lack of variation in the intonation contour. Intonation conveys, to a certain degree, the syntactic structure of an utterance and thus allows words to be grouped into meaning-carrying chunks ^26^. This slow prosodic rhythmicity is in turn tracked by the brain ^27^. Furthermore, having been shown to aid in the parsing and grouping of continuous speech ^19,28^, we expected the prosodic hierarchy and manipulations thereof to be neurally represented by the nesting of oscillations, such that PAC would be discriminable between conditions, especially at low frequencies. Finally, we expected to observe differences between synchronizer groups for both cortical entrainment and PAC ^21^. However, since the neural sensitivity of high and low synchronizers to prosodic structure is so far unexplored, we had no directed hypotheses.

## Results

### Cortical entrainment to the prosodic hierarchy

We first quantified the extent of synchronization between the speech signal and the neural signals on a single-trial basis using pairwise phase consistency (PPC) ^34^. PPC reflects how well two time series are phase-aligned. We extracted the amplitude envelopes corresponding to each prosodic level from our stimuli and filtered the EEG signals to those same frequencies (based on the average frequency bands across all stimuli; syllable/theta: 3.5-4.7 Hz, stress/high delta: 1.8-2.8 Hz, intonation/low delta: 0.6-1.1 Hz). These frequency ranges were determined acoustically based on the speech signals’ temporal modulation spectrum (see **Figure 1A**; **Methods**). We then analyzed cortical entrainment as a function of prosodic manipulations, synchronizer group, and regions of interest (ROIs) by fitting three decision tree-based mixed effects models ^35^. These iteratively partition the data into nodes based on splitting variables’ importance in explaining variance in the data, and they are suitable to detect subgroups in clustered data (such as EEG electrode sites). The root node is determined by the most significant predictor followed by splits based on other predictors, leading to terminal nodes showing (kernel density estimates of) fitted PPC values (**Figures 2-4**).

**Figure 2.**
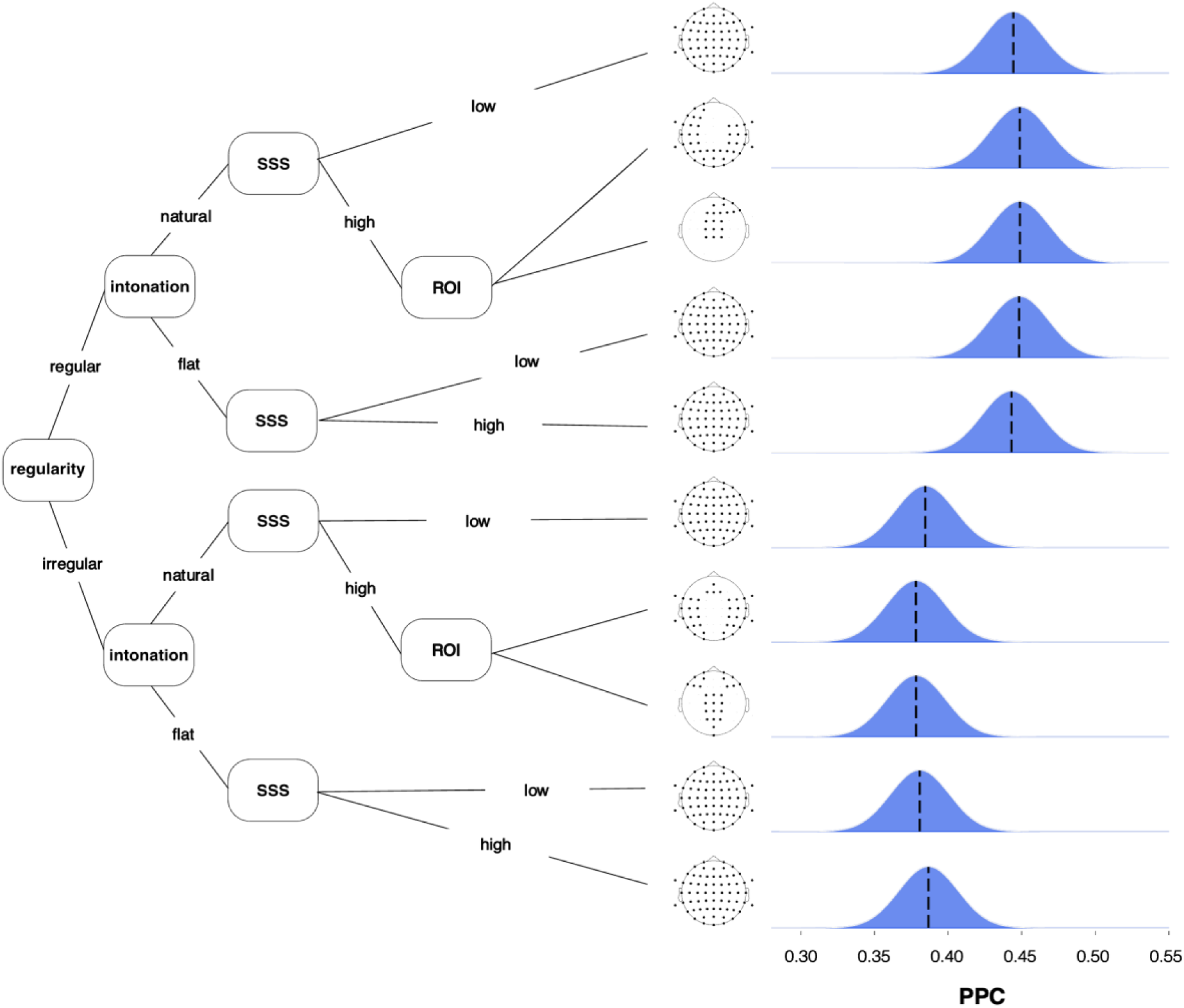
Regression model tree for pairwise phase coherence in the theta frequency range (3.5-4.7 Hz), corresponding to the syllable level in the prosodic hierarchy. Topographic maps show the electrode positions of regions of interest (ROIs) included in the respective grouping. All splits are statistically significant at *p* < .05 (Bonferroni-corrected). Terminal nodes show kernel density estimates of fitted PPC values. The dashed line represents the mean value. SSS = Spontaneous Speech Synchronization.

**Figure 3.**
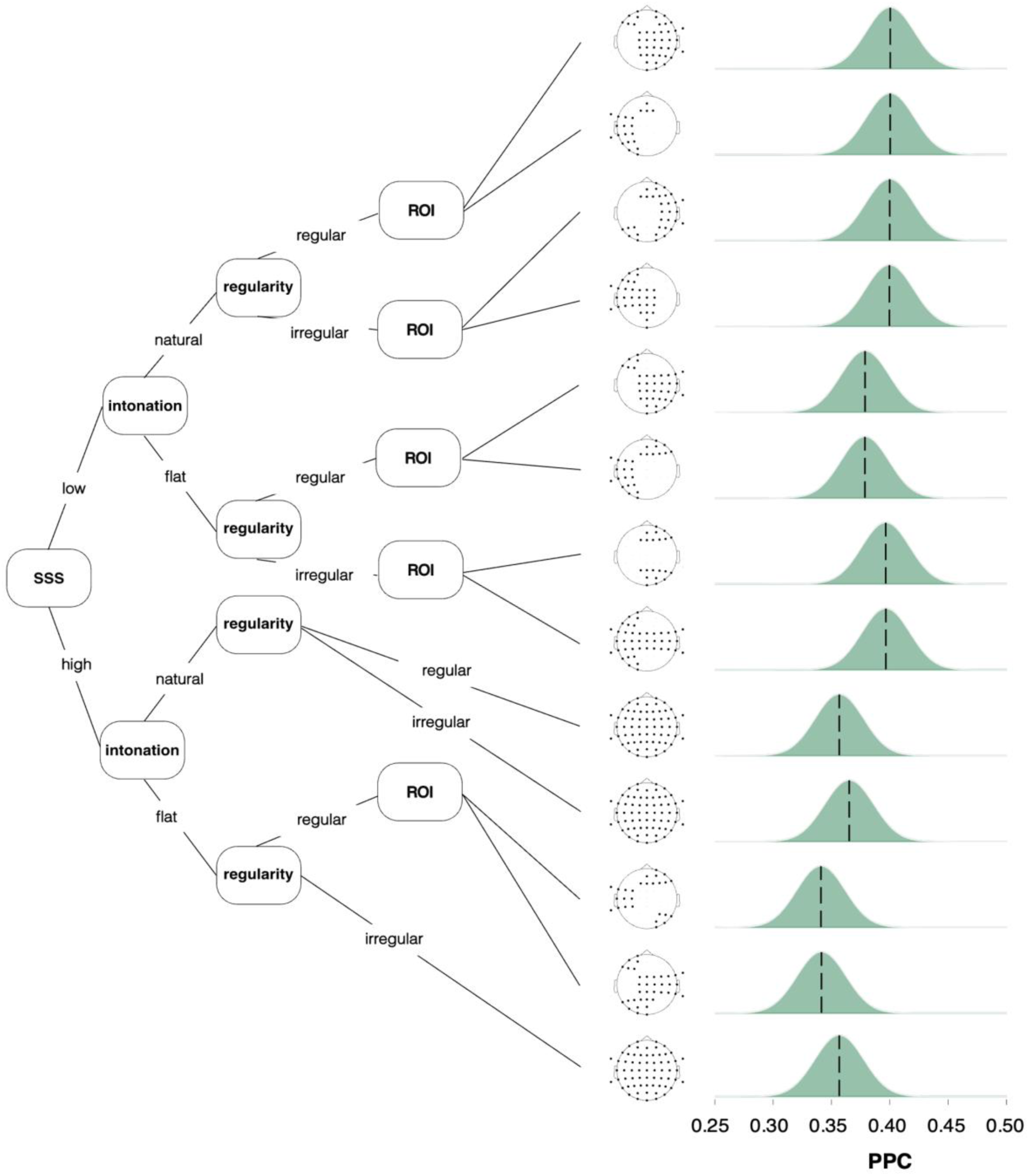
Regression model tree for pairwise phase coherence in the high delta frequency range (1.8-2.8 Hz), corresponding to the foot level in the prosodic hierarchy. Topographic maps show the electrode positions of regions of interest (ROIs) included in the respective grouping. All splits are statistically significant at *p* < .05 (Bonferroni-corrected). Terminal nodes show kernel density estimates of fitted PPC values. The dashed line represents the mean value. SSS = Spontaneous Speech Synchronization.

**Figure 4.**
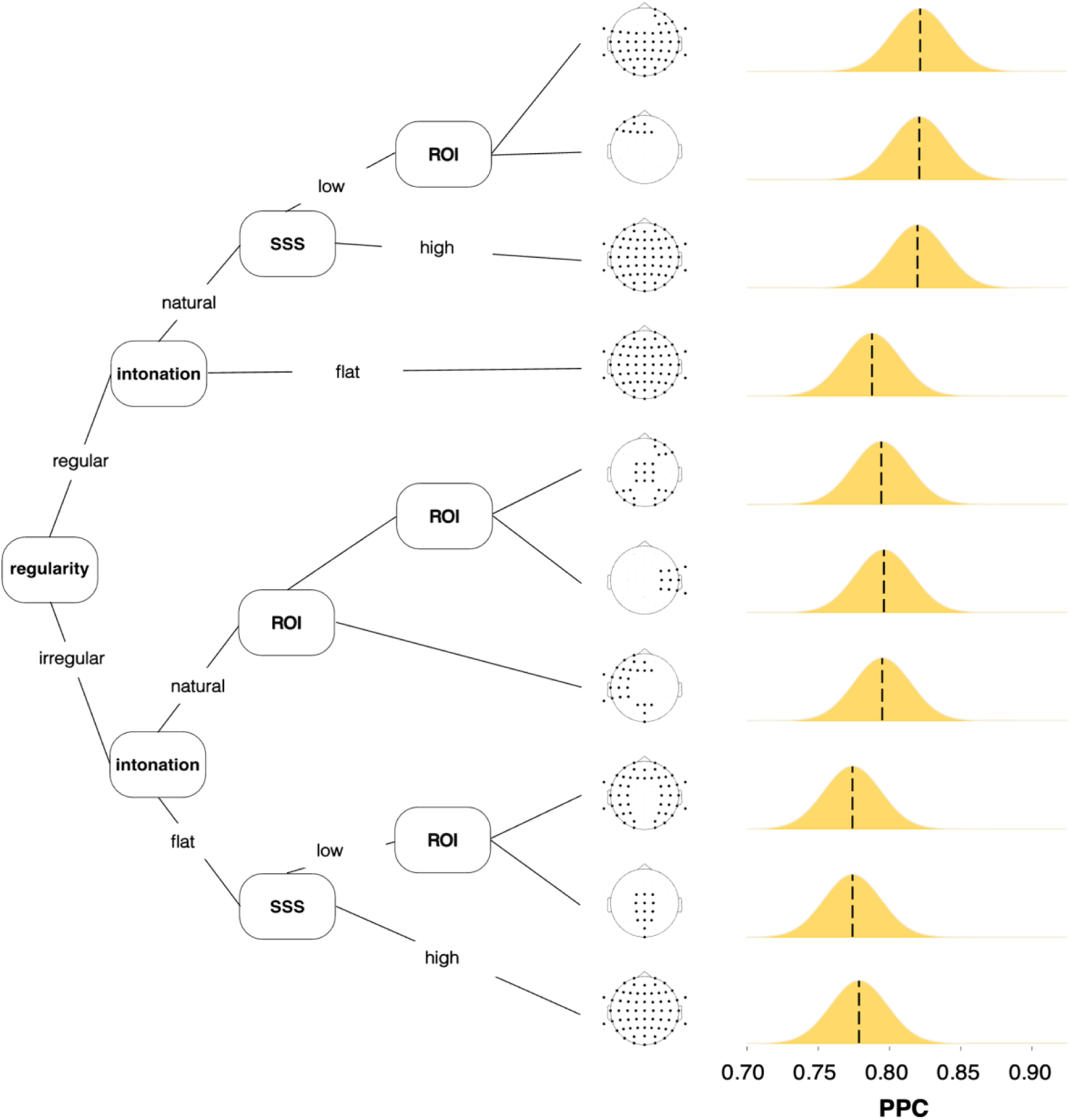
Regression model tree for pairwise phase coherence in the low delta frequency range (0.6-1.1 Hz), corresponding to the intonation phrase level in the prosodic hierarchy. Topographic maps show the electrode positions of regions of interest (ROIs) included in the respective grouping. All splits are statistically significant at *p* < .05 (Bonferroni-corrected). Terminal nodes show kernel density estimates of fitted PPC values. The dashed line represents the mean value. SSS = Spontaneous Speech Synchronization.

#### Theta/syllable

Modeling neural synchronization in the theta range (3.5-4.7 Hz) to amplitude modulations associated with syllables revealed as a first split a facilitative effect of rhythmic regularity (Figure 2). Listening to sentences with a regular stress pattern yielded higher PPC values than listening to irregularly stressed sentences. Within the regular and irregular sentences, respectively, a natural intonation contour resulted in higher neural synchronization than a flattened intonation contour. Additionally, we found (albeit somewhat varied) differences between synchronizer groups: Whereas high synchronizers exhibited stronger neural synchronization for sentences that are irregular with a flattened intonation contour and sentences that are regular with a natural intonation contour, low synchronizers showed higher PPC values for sentences that are irregular with a natural intonation contour and sentences that are regular with a flattened intonation contour. In other words, high synchronizers showed increased neural synchronization compared to low synchronizers in response to sentences that were either highly predictable (i.e., with a regular stress pattern and naturally varying intonation contour) or hard to predict (i.e., with an irregular stress pattern and flattened intonation contour). Finally, although prosodic manipulations affected neural synchronization across the whole scalp, topographical differences emerged between synchronizer groups. High synchronizers showed differences between centro-frontal and lateral-posterior electrode sites, while low synchronizers showed effects spanning the whole scalp.

#### High delta/foot

Modeling neural synchronization in the high delta range (1.8-2.8 Hz) to amplitude modulations associated with prosodic feet revealed as a first split a group difference between synchronizers (**Figure 3**). Contrary to our expectations, regardless of the particular sentence type they were listening to, low synchronizers showed increased PPC values compared to high synchronizers. Nonetheless, the next and most prominent prosodic split revealed that sentences with a flattened intonation contour elicited decreased neural synchronization in both groups. While rhythmic regularity was associated with increased PPC values in the theta frequency range, the opposite was the case for high delta, where rhythmic irregularity in both synchronizer groups and intonation conditions led to higher PPC values compared to rhythmic regularity. Finally, topographical differences emerged between rhythm conditions but especially amongst low synchronizers who mainly showed variation between electrode sites over the left and right hemispheres.

#### Low delta/intonation phrase

Modeling neural synchronization in the low delta range (0.6-1.1 Hz) to amplitude modulations associated with intonation revealed as a first split a facilitative effect of rhythmic regularity (**Figure 4**). Listening to sentences with a regular stress pattern resulted in stronger neural synchronization than listening to sentences with an irregular stress pattern. The next split revealed that a natural intonation contour is associated with higher PPC values in both regular and irregular sentences. Within sentences with a regular stress pattern and a natural intonation contour, high synchronizers exhibited effects across the whole scalp while low synchronizers showed differences between left frontal electrode sites and the rest of the scalp. Low synchronizers also showed differences between centro-parietal electrode sites and the rest of the scalp for sentences with an irregular stress pattern and a flattened intonation contour. In this condition with unpredictable stress and intonation patterns, high synchronizers again showed higher neural synchronization compared to low synchronizers whereas the opposite was true for the fully predictable condition (i.e., a regular stress pattern and naturally varying intonation contour). Finally, listening to irregular sentences with a natural intonation contour was associated with topographical differences mainly between left frontal, temporal, and posterior electrode sites and right frontal, central and posterior electrode sites.

Overall, the results from all three models imply that the amplitude modulations associated with the levels of the prosodic hierarchy modulate the brain’s response in the form of top-down cortical entrainment (**Figure 5**). Acoustic manipulations at higher levels in the prosodic hierarchy mainly influenced cortical entrainment at lower levels, implying a hierarchy not only in the acoustic and phonological space but also embodied in neural activity. Moreover, the results indicate that listeners are differentially sensitive to the speech signal’s prosodic properties and accordingly vary in their neural responses to it.

**Figure 5.**
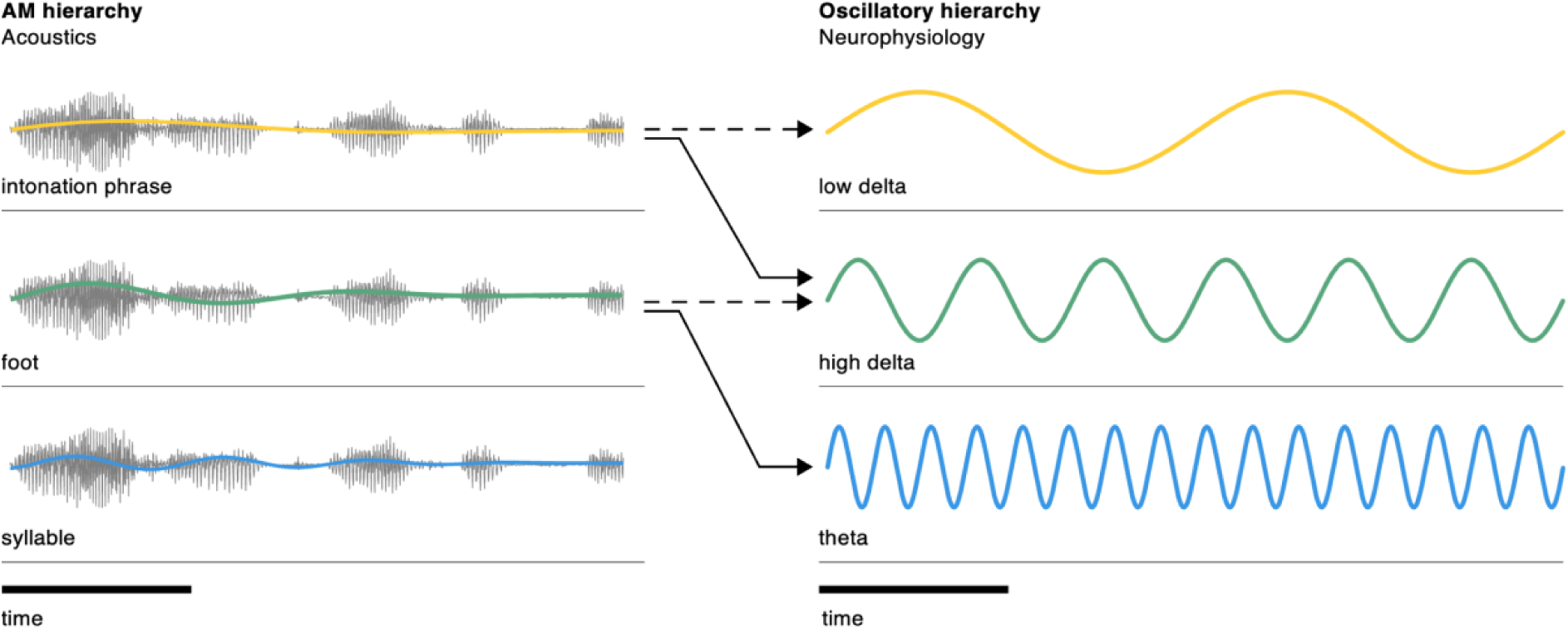
Taking into account the results from all three regression models reveal that the acoustic information associated with the prosodic hierarchy top-down modulates cortical entrainment. As we illustratively show in this overview figure, the main influence on a given level originates from prosodic manipulations at the next higher, dominating level (bold lines). Influence from manipulations at the corresponding level are also present, albeit weaker (dashed lines).

### Decoding of phase-amplitude coupling

To further explore this pattern of hierarchical relationships, we tested the extent to which information about the speech signal is represented neurally by the nesting of oscillations (see **Methods**). We first calculated phase-amplitude coupling using non-linear driven autoregressive models ^36^. The resulting comodulograms reflect the strength of coupling between the phase of a low-frequency driver and the amplitude of a higher signal frequency. We further reasoned that if the speech signal has distinct neural representations between conditions in the form of PAC, the corresponding comodulograms should be reliably discriminable by a classifier ^37^. We therefore performed decoding by training a linear support vector machine (SVM) to discriminate between experimental conditions and their associated PAC comodulograms. Additionally, while achieving above-chance-level decoding accuracy suggests that classification is not purely random, these thresholds really only hold for infinite sample sizes ^38^. Therefore, we only report decoding accuracies that we found to be significant (FDR-corrected at *p* < .05) using surrogates and permutation testing (see **Methods**).

Following these steps, **Figure 6A-C** shows that across the scalp, there are electrodes that reflect neurally distinct PAC between conditions and synchronizer groups. The SVM classifier was able to distinguish whether PAC resulted from trials with a regular vs. irregular rhythm, a natural vs. flattened intonation contour, or whether the comodulogram stemmed from a high or a low synchronizer. Significant electrodes for regularity (**Figure 6A**) and synchronizer group (**Figure 6C**) were more numerous over the left hemisphere than over the right hemisphere, whereas they were more distributed across the whole scalp for intonation (**Figure 6B**). In an 8-class classification, the classifier was also able to distinguish the combination of prosodic manipulations and synchronizer group in a fine-grained manner (**Figure 6D**). Electrodes that showed significant above chance level decoding accuracies were distributed across the whole scalp. These results suggest that how oscillations are nested depends on the prosodic hierarchy and manipulations thereof, as well as on synchronizer group.

**Figure 6.**
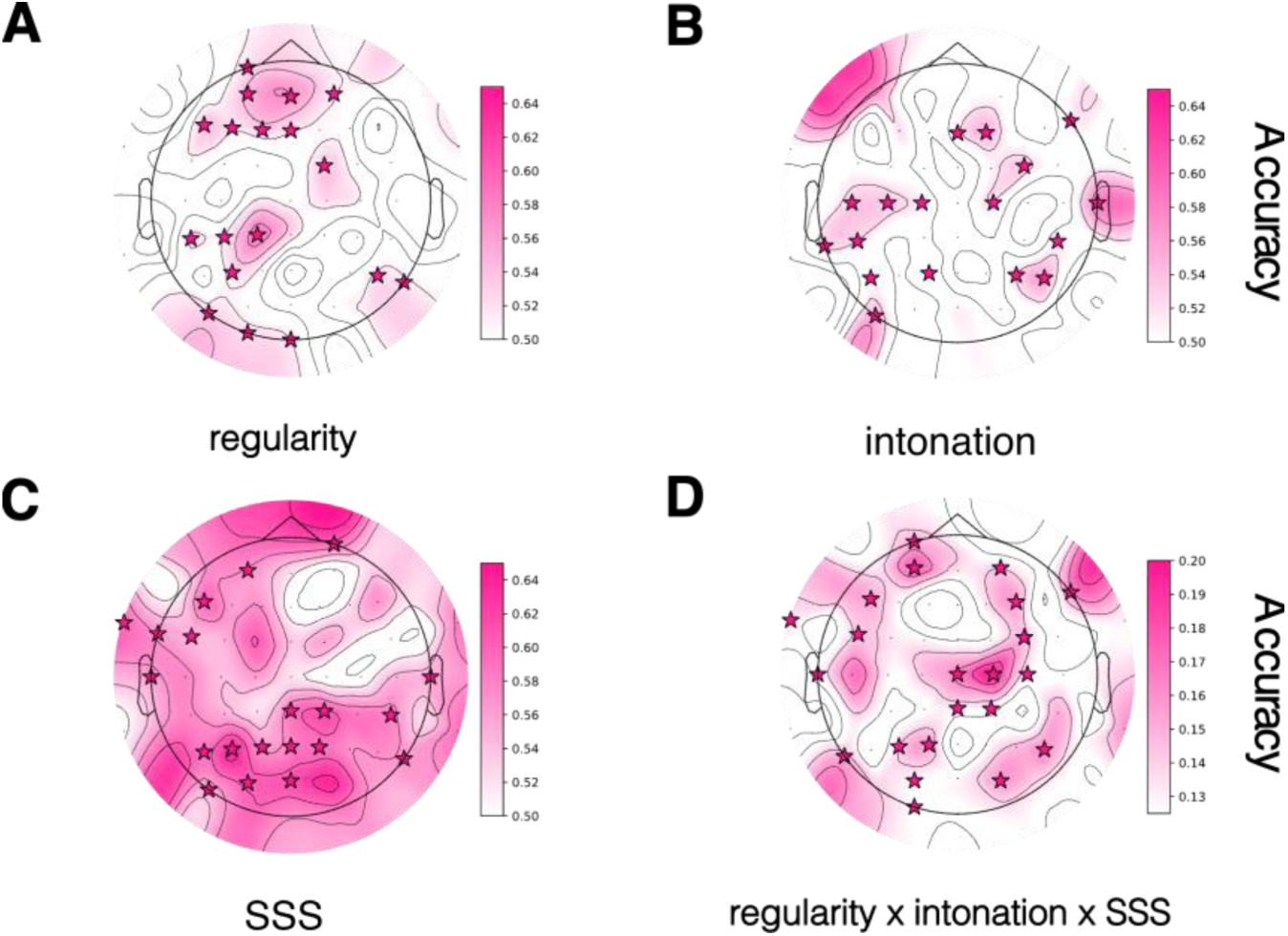
Topographic plots of decoding accuracies. Stars denote electrodes identified by permutation testing as showing above-chance-level decoding accuracies significantly different from decoding accuracies stemming from classifying surrogate PAC comodulograms. **A** Decoding accuracies achieved by the classifier when distinguishing comodulograms stemming from regular vs. irregular sentences (chance level = 50%). **B** Decoding accuracies achieved by the classifier when distinguishing comodulograms stemming from sentences with a naturally varying vs. flattened intonation contour (chance level = 50%) **C** Decoding accuracies achieved by the classifier when distinguishing PAC comodulograms stemming from high vs. low synchronizers (chance level = 50%). **D** Decoding accuracies achieved by the classifier when distinguishing fully crossed PAC comodulograms (i.e., 8 classes; chance level = 12.5%). SSS = Spontaneous Speech Synchronization.

To identify PAC between which frequencies drove classification, we extracted inverse-transformed support vector weights from the fitted classifiers. With their magnitude and direction, they indicate the importance of the contribution of each feature to the decision boundary and thus the classification performance. To enable a neurophysiological interpretation, we extracted the weights as spatial patterns ^39^. Plotting these patterns (averaged across electrodes showing significant above-chance-level decoding accuracy) shows that nesting between the phase of oscillations between ∼0.5 and 4 Hz and the amplitude of low-frequency oscillations between ∼1.5 and 8 Hz mainly acted as drivers of classification of prosodic condition as well as synchronizer group (**Figure 7**). Also important for classification, albeit to a lesser extent, was PAC between the phase of oscillations between ∼0.5 and 10 Hz and the amplitude of higher-frequency oscillations around 30 and 45 Hz. On the one hand, this is evidence that the speech signal’s hierarchical information occurring at low modulation frequencies is dealt with by oscillatory cross-frequency interactions. On the other hand, this also shows that these cross-frequency interactions manifest differently depending on whether a person is a high or a low synchronizer.

**Figure 7.**
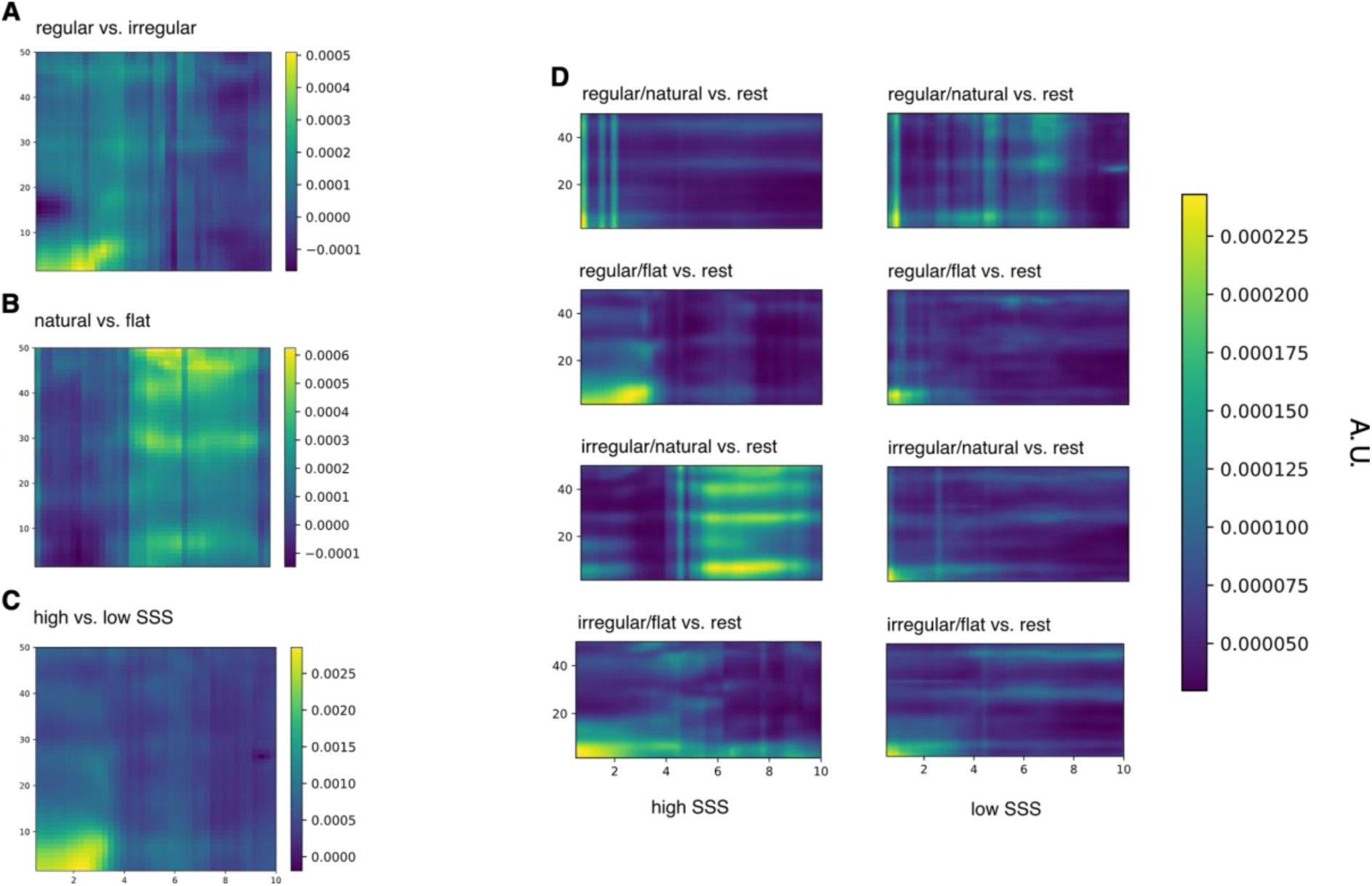
Matrices of inverse-transformed classifier weights extracted as spatial patterns, indicating how much a particular feature contributed towards the classifier’s decision. The main drivers of classification were PAC between the phase of oscillations between ∼0.5 and 4 Hz and the amplitude of oscillations between ∼1.5 and 8 Hz. Additionally, PAC between the phase of oscillations between ∼0.5 and 10 Hz and the amplitude of higher-frequency oscillations ∼30 and 45 Hz also contributed to the classification decision. **A** Weights indicating which features were driving the classification of PAC comodulograms stemming from trials with a regular vs. irregular rhythm. **B** Weights indicating which features were driving the classification of PAC comodulograms stemming from trials with a naturally varying vs. flattened intonation contour. **C** Weights indicating which features were driving the classification of PAC comodulograms stemming from high vs. low synchronizers. **D** Weights indicating which features were driving classification of fully crossed PAC comodulograms.

## Discussion

This study brings into alignment acoustics, phonology, and neurophysiology to probe the processing of spoken language in the brain. The combined cortical entrainment and phase-amplitude coupling responses to the syllable, foot, and intonation phrase levels invite the conclusion that the prosodic hierarchy has a specific neural implementation. Evaluating the extent to which the brain synchronizes to the speech signal showed that listeners concurrently process the amplitude modulations associated with the prosodic hierarchy. We found that this neural response is structured in such a way that reflects this hierarchy: Amplitude modulations associated with a given level of the prosodic hierarchy primarily influenced neural synchronization of subordinate levels. Regular and irregular alternations of stress at the foot level had the greatest impact on cortical entrainment at the syllable level (i.e., in the theta range (3.5-4.7 Hz)). Similarly, manipulating the intonation contour had the greatest influence on cortical entrainment at the foot level (i.e., the high delta range). In both instances, we observed a facilitative effect when the speech signal contained predictable prosodic rhythms, i.e., a regular (vs. an irregular) stress pattern or a naturally varying (vs. flattened) intonation contour. We suggest that underlying these effects are the predictive mechanisms inherent to entrainment. In synchronizing to the regularities of the speech signal, oscillatory cycles act as temporal windows to chunk discrete linguistic units ^14,15^, in our case syllables, feet, and intonation phrases. It is assumed that the phases of these cycles can be used by brain areas further down in the processing stream (e.g., the motor system or perisylvian areas of higher-level linguistic processing) to predict the occurrence of incoming auditory events ^40,41^. Top-down predictions stemming from these areas phase-reset cortical oscillations ^40^ such that information coinciding with phases of high neuronal excitability are, as a consequence, amplified and easier to perceive ^42^, ultimately giving rise to a facilitative effect.

Furthermore, delta-theta PAC varies as a function of the prosodic hierarchy and the manipulation thereof. Hierarchically coupled low-frequency oscillations (< 8 Hz) were the main features that differentiated PAC comodulograms. This suggests that the brain employs nested oscillations to concurrently process and represent the acoustic information associated with the prosodic hierarchy. Doing so presumably allows the brain to maintain this hierarchy and the co-dependent nature of the levels ^14,18,28,43^. As a consequence, emerging from this top-down coordination of neural activity are discrete units where syllables are bound inside feet and feet are bound inside intonation phrases. The finding that PAC comodulograms were distinguishable as a function of experimental conditions furthermore indicates that introducing prosodic manipulations to the speech signal led to altered cross-frequency coupling. Depending on the intactness of the prosodic hierarchy, this would lead to (im)precisely coupled oscillations and thus (in)accurately chunked units. Although this has otherwise not been tested in the context of prosody, previous research has shown that neural activity in the form of phase-amplitude coupling is less coordinated for non-speech ^44^, unintelligible speech ^28,45^, and non-native speech ^45^. These findings show that the robustness of PAC coordination is related to the efficiency with which the brain can chunk the speech signal. However, it remains to be seen whether the degree of phase-amplitude coupling as a function of the prosodic hierarchy is also related to behavior or perception and thus is truly conducive to efficient parsing. We additionally found that theta-gamma coupling likewise (although to a lesser extent) contributed to PAC comodulogram classification. Since we did not systematically vary acoustic information in the gamma frequency range (∼30/45 Hz), this activity could be indicative of “the operating rhythm of the sensory cortex” ^18^ related to the (de-)activation of neuronal networks ^17^ that varied between experimental conditions. In instances where a stimulus contains predictable rhythmic patterns, the brain operates in a rhythmic mode where lower frequency phase and higher frequency (gamma) amplitude are coupled. Gamma oscillations become more synchronous whenever a task-relevant event is expected ^46^, enhancing neural representations. When rhythmic patterns are absent or unpredictable, the operating mode changes to one of continuity, and oscillations become decoupled ^46^. Although these conjectures in the context of our study are speculative for the time being, these different operating modes may be what partially drove classification.

The burgeoning of research on and theorizing about neural oscillations notwithstanding, there is an ongoing debate about the nature of entrainment and whether it is to be considered epiphenomenal, a methodological artifact, or rather fundamentally causal in the context of language and cognition ^47^. Contributing to this discussion, the top-down, hierarchical structure of low-frequency neural responses to the prosodic hierarchy we find here speaks to the third interpretation. Low-frequency (delta-band) oscillations, often found in associative ^48^ and frontal areas ^49,50^, have been shown to top-down modulate both gamma power ^49^ as well as entrainment ^50^ in (left) auditory cortex, presumably harnessing contextual information like prosody ^3,15^ and conveying predictions from higher to lower levels ^51^. Their observed role in serving linguistic and speech-specific processing ^52,53^ therefore comes as no surprise. Additionally, neural phase synchronization in the three frequency bands showed non-redundant effects. If the results were merely reflective of passive bottom-up tracking of acoustic transients, we would have expected to observe identical effects across all three frequency bands and no top-down modulation ^54^. Finally, it has been argued that it is difficult to distinguish the spectro-temporal patterns of evoked responses from those of genuine oscillations ^55^. However, in the case of low-frequency entrainment, evoked responses like the N100 cannot make any measurable contribution on account of their short latencies while long-latency components like the N400 or P600, although plausibly involved, are unlikely to have differed systematically between our conditions since semantic and syntactic properties were controlled for ^56^.

Language and speech are such integral parts of the human condition that - barring pathology - most humans can be considered experts. At the same time, variation is a natural consequence of the dynamic and complex processes of life that are expressed in our genetic makeup and behaviors, including language and speech. This does not just pertain to variation as found in disorders like specific language impairment, aphasia, or autism but, as it turns out, also to more subtle differences in typically-developing populations. Recently, Assaneo et al. ^21^ uncovered a bimodal distribution in the population: High and low synchronizers show systematic differences with respect to brain anatomy, electrophysiological function, and behavior. The behavioral, audio-motor synchronization task they developed to identify these groups requires listeners to synchronize their own speech output with a syllable stream. This task mainly targets the rhythmic amplitude modulations associated with syllabic information and captures the extent to which individuals can capitalize on efferent motor signals to predict auditory input ^21^. However, as shown in **Figure 1A**, the speech signal also contains other prominent prosodic rhythms. So far, it remains unexplored whether auditory-motor coupling as measured by this task also relates to and is beneficial for the perception of these slower rhythms as a function of group membership. While we did find some group differences regarding cortical entrainment, these varied across frequency ranges and showed that high synchronizers are not pervasively more in tune with the rhythmicity of the speech signal (although they tended to show stronger synchronization to conditions with unpredictable rhythms). These varying patterns of cortical entrainment might come about as a consequence of compensatory mechanisms, non-linear relationships, or differential allocation of resources ^57^. Despite these discrepancies, decoding of PAC patterns revealed more systematic differences in the neural representations of the prosodic hierarchy between the two groups especially in CFC between low-frequency oscillations. We propose that at the heart of these findings lie qualitative differences in how the motor system is engaged during perception such that the top-down phase-reset of oscillations elicited by the motor system results in more or less precisely aligned high excitability phases with the incoming sensory input ^40,58,59^. Although the results differentiating synchronizer groups vary across neural measures and experimental conditions, they are nonetheless compelling and imply that the differential engagement of the motor system has an effect on the representation of prosodic information. Nevertheless, the group differences we found here as well as the involvement of the motor system in prosody perception need to be explored further.

We provide evidence that prosody and its hierarchical structure are cues that decisively shape the neural representation of the speech signal and the efficiency with which this representation is formed. By interfacing acoustics, phonology, and neurophysiology, we characterize one of the complex ways in which the rich information in the speech signal is elaborated by the brain. The experiment underscores the involvement of predictive oscillatory mechanisms in this process, and it extends the audio-motor synchronization framework to include prosodic rhythms, thus implying the motor system as a pivotal mediator of prosodic perception. Finally, our results substantiate the assumptions brought to bear in theories about prosodic structure, revealing that the proclaimed hierarchy in prosody is maintained at a neural level. Whether this holds universally across languages will need to be explored in future research. Overall, this set of findings informs and broadens current theories on spoken language comprehension in the brain as well as phonological research.

## Methods

### Participants

We recruited 33 native speakers of Swiss German (5 males). Participants were right-handed, non-musicians and none reported any past or current neurological or severe psychiatric disorders. Three participants were excluded from the study: We had technical issues with one, another was not a native speaker of Swiss German and the last was not right-handed. This left us with a sample size of 30 participants between the ages of 19 and 29 years (M = 23.2 years, SD = 3.32 years). We assessed and confirmed normal hearing ability using pure tone audiometry (MAICO ST20, MAICO Diagnostics, Berlin, Germany), defined as audiometric thresholds of 30 dB HL at octave frequencies from 0.25 to 8.0 kHz. We determined participants’ handedness using a German version of the Annett questionnaire ^60^. Participants were informed in written and verbal form of the procedure and gave their written consent. Participants received either course credit or monetary compensation for their efforts. The study protocol was approved by the ethics committee of the Faculty of Arts and Social Sciences, University of Zurich (Approval number: 20.6.12), and conducted according to the principles defined in the Declaration of Helsinki.

### Stimuli

The stimulus material consisted of 480 unique German sentences spoken by a trained, female native speaker who was instructed to speak naturally and clearly. The recorded sentences were between 3403 and 4505 ms long (M = 3954 ms, SD = 177 ms). Sentences followed a constant syntactic structure: a noun phrase (either a proper name or a noun), a verb phrase (a modal verb modified by an adverbial phrase), and an object noun phrase (modified by one or two adjectives). Sentences were constructed to be meaningful but low in semantic predictability such that participants would need to attend to and precisely analyze each element. All words were part of the basic German vocabulary ^61^ (among the 10000 most frequent words). To exacerbate the brain’s ability to concurrently represent different levels of the prosodic hierarchy, any given sentence always consisted of 16 syllables and contained experimental manipulations at the levels of the foot and intonation phrase. At the level of the foot, sentences either followed a predictable regular (with consistently alternating stressed and unstressed syllables) or an unpredictable irregular (but still legal) stress pattern (e.g., regular: Christa sollte damals einen runden gelben Käfer malen; irregular: Leonardo soll anschliessend den berühmten Hut aufsetzen). There were two further conditions whose stress pattern changed after 8 syllables from regular to irregular or irregular to regular, respectively. For the purposes of this study, we restricted our analyses to the first 8 syllables. At the level of the intonation phrase, sentences either had a naturally varying or a constant intonation contour. We used Praat ^62^ to extract the mean fundamental frequency (f0) value across all stimuli (196.61 Hz) and flatten individual sentence’s f0 to this value with the Pitch-Synchronous Overlap-Add (PSOLA) algorithm. Stimuli were presented at a sampling rate of 44100 Hz.

### Procedure

From the pool of 480 stimuli, we randomly assigned 400 spoken sentences to each participant. Half of these sentences were then randomly assigned to the pitch-flattened intonation condition and the other half to the natural intonation condition. This means that a given participant never heard the same stimulus in both intonation conditions. During EEG recording, participants were comfortably seated inside an electromagnetically-shielded booth. They were binaurally presented with the stimuli through insert earphones (Etymotic ER-2, Etymotic Research) attached to foam ear tips (ER1-14A) in a randomized order in five blocks of 80 trials at a volume of approximately 70 dB SPL. We used Presentation (Neurobehavioral Systems Inc., Berkeley, USA) to control stimulus delivery. Participants took short breaks between blocks as needed. They first completed a series of practice trials. As a behavioral measure, participants were asked to type on a keyboard what had been said in the noise after listening to each stimulus. There was no time limitation for participants’ responses, but they were encouraged to write their responses spontaneously and even if they had not understood every single word. The inter-trial interval following this response was randomly jittered between 2000 and 2500 ms. The interval between offset of speech and onset of response screen was jittered between 300 and 400 ms. The main experiment (excluding EEG application) lasted approximately 80 minutes.

### Speech preprocessing

For each stimulus, we computed the broadband envelope in Python, using the toolboxes Brian2 (version 2.4.2), Brian2Hears (version 0.9.2) ^63,64^ and Scipy (version 1.5.3). First, to simulate the basilar membrane within the cochlea, we applied a gammatone filterbank ^1,65^: Acoustic waveforms were filtered into 8 subbands in the human hearing range from 20 Hz to 20’000 Hz and center frequencies equally spaced on the approximately logarithmic equivalent rectangular bandwidth (ERB) scale. Gammatone filters were implemented as cascades of four 2^nd^ order IIR filters. The approximated impulse response is based on the exact gammatone implementation as described by Slaney ^66^. The parameter values for computation of the ERB bandwidth stem from Glasberg and Moore ^67^. To include further non-linearities of cochlear processing – specifically the transduction properties of inner hair cells – we then applied half-wave rectification and compression with a 6/10 power law to the output of the gammatone filterbank. We recombined the subbands as the average to obtain the broadband envelope. Finally, the signal was downsampled from 44100 Hz to 1000 Hz. The continuous f0 envelope was calculated by first extracting the raw pitch values for each stimulus using Praat ^62^ and then interpolating these using the Wavelet prosody analyzer toolkit (version 3.2.2020) ^68^. The frequency bands of interest for which to compute phase synchronization were then determined based on the peaks in the temporal modulation spectra of the speech stimuli (**Figure 1A**; see below) ^69,70^. These acoustically defined ranges corresponded to phonologically defined ranges of the syllable, stress, intonation phrase rates (i.e. the number of produced syllables, stressed syllables/feet, and intonation phrases divided by sentence duration). The minimum and maximum across all these values determined the lower and upper bound of the respective frequency bands: syllable/theta: 3.5-4.7 Hz, stress/high delta: 1.8-2.8 Hz, intonation/low delta: 0.6-1.1 Hz. The envelopes were then filtered to these frequencies using a 3^rd^ order Butterworth bandpass filter (roll-off 18 dB/octave; forward-backward digital filter using cascaded second-order sections).

### Temporal modulation spectrum

The temporal modulation spectrum of a sound signal shows how fast its amplitude fluctuates over time and is neurophysiologically interpreted to approximately reflect the tuning of neurons to modulation frequencies ^69,71,72^. For its calculation, we followed the procedure described in Ding et al. ^69^. First, for each speech stimulus we extracted its amplitude envelope (as described above). Then each envelope’s discrete Fourier Transform (DFT) is computed by means of the Fast Fourier Transform (FFT). The modulation spectrum is equal to the root-mean-square of the DFT of all envelopes. We ran these calculations for speech stimuli with a natural vs. a flat intonation contour separately (see **Figure 1A**).

### EEG

EEG was recorded using BrainVision Recorder (version 1.23.0003, BrainProducts GmbH, Gilching, Germany) and a 64-channel actiCHamp Plus system (Brain Products GmbH, Gilching, Germany) arranged according to an extended 10/20 system with the following Ag/AgCl active electrodes: Fp1, Fp2, AF7, AF3, AFz, AF4, AF8, F7, F5, F3, F1, Fz, F2, F4, F6, F8, FT9, FT7, FC5, FC3, FC1, FC2, FC4, FC6, FT8, FT10, T7, C5, C3, C1, Cz, C2, C4, C6, T8, TP9, TP7, CP5, CP3, CP1, CPz, CP2, CP4, CP6, TP8, TP10, P7, P5, P3, P1, Pz, P2, P4, P6, P8, PO7, PO3, POz, PO4, PO8, O1, Oz, O2. The signal was online referenced to FCz and we used electrode FPz as ground. Throughout the recording, electrode impedance was kept under 30 kΩ using electroconductive gel. The signal was recorded continuously with a bandwidth from DC to 280 Hz and digitized at a sampling rate of 1000 Hz. Preprocessing of the data was performed in Python (version 3.8.6) using MNE Python (version 1.0.0) ^73,74^. We first filtered the data from 1 Hz to 30 Hz (for phase synchronization analyses) and from 1 Hz to 50 Hz (for phase-amplitude coupling analyses) using a one-pass, zero-phase, non-causal bandpass filter (roll-off 6 dB/octave) and applied a one-pass, zero-phase non-causal bandstop filter at 50 Hz and harmonics (roll-off 6 dB/octave). Ocular artifacts such as blinks and saccades and heartbeat artifacts were corrected using independent component analysis (ICA) ^75^. They were manually selected by visually inspecting components’ topographies and time courses. Artifactual channels were identified visually and excluded from the ICA decomposition. The ICA solution was then applied to data filtered between 0.3 Hz and 30 Hz (and 0.3 Hz and 50 Hz) (one-pass, zero-phase, non-causal bandpass filter; roll-off 6 dB/octave) and previously excluded channels were interpolated using spherical spline interpolation ^76^. We re-referenced the cleaned data to the average across all channels and segmented the continuous data into epochs from –2000 ms prior to speech onset to 300 ms post speech offset. Finally, epochs with a maximum and/or minimum peak-to-peak signal amplitude (PTP) of 150 and 1µV, respectively, were dropped. PTP is defined as the absolute difference between the lowest and highest signal value and indicates the peakedness/flatness of the signal.

### Phase synchronization

EEG signals were first bandpass filtered to the same frequencies as the speech signal (3.5-4.7 Hz, 1.8-2.8 Hz, and 0.6-1.1 Hz, respectively) using a 3^rd^ order Butterworth bandpass filter (roll-off 18 dB/octave; forward-backward digital filter using cascaded second-order sections). For each signal, we computed the Hilbert transform and extracted instantaneous phase as the phase angle of the resulting analytic signal. To quantify the extent of phase synchronization, we calculated single-trial pairwise phase consistency (PPC) ^34^ between each speech stimulus and each EEG channel. An advantage of the PPC is that it is not biased by sample size. However, it is computationally less efficient than other measures of phase synchrony ^77^. To counter this, we further downsampled all involved signals to 100 Hz. We omitted the first 500 ms of the sentences from this calculation to avoid spurious phase-locking induced by evoked potentials after speech onset. PPC is calculated as follows: First, the relative phase between the EEG and speech signal at each timepoint within a trial is calculated. Then, the angular distance between each observation of relative phase is calculated pairwise. If the phase relationship between the two signals is synchronous across time, the angular distances between pairs of relative phases will be small. Finally, averaging the dot product across all pairs of these observations results in the PPC. PPC values of 0 indicate non-existent phase synchronization while values of 1 indicate perfect phase synchronization.

### Cross-frequency coupling

To test for phase-amplitude coupling (PAC) and whether it varies as a function of experimental manipulations, we computed comodulograms on the basis of non-linear driven autoregressive (DAR) models using the Python toolbox pactools (version: 0.3.1) ^36^. This is a statistical signal modeling approach (as opposed to a biophysical one) and it avoids some of the methodological pitfalls associated with other PAC measures such as applying the Hilbert transform to broadband signals or incorrectly choosing the bandwidth of bandpass filters. DAR models can handle the non-linearity and non-stationarity characteristic of neural signals and do not expect these to be strictly sinusoidal. Specifically, the (low) driver frequency is first extracted using a bandpass filter. The high frequencies result from subtracting this driver from the raw signal thus modeling them globally. The resulting gap in the signal is filled with Gaussian white noise. By repeating this procedure for a range of driver frequencies, we obtain a comodulogram that reflects PAC between pairs of frequencies.

Applied to our data, signals were first downsampled to 250 Hz. For each participant and electrode-condition combination, a comodulogram was then calculated across trials for low frequencies between 0.5 and 10 Hz and high frequencies between 1.5 and 50 Hz using an autoregressive model order of 20. In the absence of actual phase-amplitude coupling, a measure of PAC can yield values close to but not exactly zero. Thus, to distinguish spurious from actual PAC, we created surrogate comodulograms by adding random time shifts, thereby removing any possible coupling ^36^.

### SSS test

To allow a classification into high and low synchronizers, participants completed the Spontaneous Speech Synchronization (SSS) task ^21^ where they hear a stream of syllables and are asked to whisper the syllable “ta” simultaneously and continuously. Analysis of the degree of synchronization between the produced and perceived speech stream and the categorization into high and low synchronization was done using the openly available code ^22^. This resulted in 11 high and 19 low synchronizers (high synchronizers: M _PLV_ = 0.59, SD _PLV_ = 0.05; low synchronizers: M _PLV_ = 0.28, SD _PLV_ = 0.06).

### Statistical analyses

#### Regions of interest

For statistical analysis of phase synchronization, channels were grouped into nine regions of interest (ROIs): left frontal (Fp1, AF7, AF3, F7, F5, F3), middle frontal (AFz, F1, Fz, F2), right frontal (Fp2, AF4, AF8, F4, F6, F8), left central (FT9, FT7, FC5, FC3, T7, C5, C3, TP9, TP7, CP5, CP3), middle central (FC1, FCz, FC2, C1, Cz, CP1, CPz, CP2), right central (FC4, FC6, FT8, FT10, C4, C6, T8, CP4, CP6, TP8, TP10), left posterior (P7, P5, P3, PO7, PO3, O1), middle posterior (P1, Pz, P2, POz, Oz), right posterior (P4, P6, P8, PO4, PO8, O2).

#### Mixed models: glmertrees

For the statistical analysis of phase synchronization, we used the R package glmertree ^78^ which implements a decision tree-based algorithm for fitting mixed effects models. Predictors (fixed and random effects), their higher-order effects and interactions are used to split the data into subgroups based on the similarity of response patterns. The model is fit to each subgroup such that different effects can be estimated for different subgroups. The relationship between dependent and independent variables are described by a binary decision tree, which has the advantage of increased interpretability ^35^. The splitting variable at each node is selected based on the p value of an association test. The value that is used to split the resulting two new nodes is chosen by optimizing the sum of their loss function^35^.

We fit a beta regression (logit link function) for each of the three frequency bands separately with regularity (regular/irregular), intonation (natural/flat), SSS (high/low), and ROI (9 ROIs) as partitioning variables. The maximal random effects structure ^79,80^ consisted of by-participant and by-item random slopes and was simplified if there were convergence or singular fit issues ^80^. For the sake of interpretability, the maximal number of splits in each model was restricted to five.

#### Multivariate-pattern analysis (MVPA) on PAC comodulograms

With the goal of investigating the extent to which information about the speech signal is represented in phase-amplitude coupling (PAC), we performed group-level decoding of PAC comodulograms across the scalp. We trained a machine learning classifier to distinguish between experimental conditions and synchronizer groups and their associated PAC comodulograms at each electrode. The resulting decoding accuracy is indicative of the information available in phase-amplitude coupling about the discriminability of experimental conditions and their respective neural representations ^37^ at the sensor-level.

The data consisted of 2D frequency-frequency matrices per electrode, per condition, and per participant. These were unravelled to vectors and associated with their respective target label (regularity: regular, irregular; intonation: natural, flat; regularity x intonation: regular/natural, regular/flat, irregular/natural, irregular/flat, regularity x intonation x SSS: regular/natural/high, regular/natural/low, regular/flat/high, regular/flat/low, irregular/natural/high, irregular/natural/low, irregular/flat/high, irregular/flat/low). We performed decoding of comodulograms using scikit-learn (version 1.2.0) ^81^ in Python (version 3.10.8). After z-transforming the data, we classified PAC comodulograms using a linear Support Vector Machine (regularization parameter C = 1). We assessed decoding accuracy with a repeated stratified 5-fold cross-validation scheme (100 iterations). This means that, for every iteration, the data was split into five roughly equal-sized folds while ensuring that each fold contained a proportion of target labels representative of the whole dataset. Classifier training was done on four folds and testing on the remaining fold. This was repeated until each fold had been used for testing once. Decoding accuracy corresponded to the mean correct classification across all folds and iterations. In the case of multiclass classification, a one-vs.-rest strategy was used.

Although decoding accuracy above chance level indicates that classification is not purely random, these levels may have limited informative value with datasets of small sample size ^38^. We thus provide statistical significance levels for decoding accuracy by relying on an empirical solution, i.e., permutation testing. Specifically, we test whether decoding accuracies of actual PAC comodulograms are significantly higher than decoding accuracies obtained from classifying surrogate PAC comodulograms (10000 permutations). **Figure 6** shows electrodes that had above-chance-level decoding in combination with being significantly different from surrogate decoding accuracies (*p* < .05, False Discovery Rate-corrected ^82^ for 64 tests).

Next, in order to gauge which features (i.e., PAC frequency pairs) drove decoding accuracy, we extracted the support vector weights associated with each feature. These weights reflect how much of a contribution a particular feature made towards the classification decision and thus to the discriminative information used by the classifier. To facilitate a neurophysiological interpretation, we extracted these weights as spatial patterns ^39^. They were inverse-transformed according to the transformer steps described above.

## Data and code availability

OSF repository: http://doi.org/10.17605/OSF.IO/82NXU

## Acknowledgements

We thank David Poeppel and Jonas Obleser for comments on earlier versions of this work, and Elisabeth Stark, Johannes Graën, and the Linguistic Research Infrastructure (LiRI) Zürich for input on the stimulus material. We also thank Marjolein Fokkema for statistical advice. This work was funded by the NCCR Evolving Language, Swiss National Science Foundation Agreement #51NF40_180888 (M.M.); S.S. is supported by Swiss National Science Foundation grant PZ00P1_208915.

## CRediT author statement

**Chantal Oderbolz:** Conceptualization, Methodology, Formal analysis, Investigation, Writing - Original Draft, Writing - Review & Editing, Visualization, Project administration. **Sebastian Sauppe:** Conceptualization, Methodology, Writing - Review & Editing. **Martin Meyer:** Conceptualization, Writing - Review & Editing, Supervision, Resources, Funding acquisition.

